# Cryo-electron microscopy structure of the SADS-CoV spike glycoprotein provides insights into an evolution of unique coronavirus spike proteins

**DOI:** 10.1101/2020.03.04.976258

**Authors:** Hongxin Guan, Youwang Wang, Abdullah F.U.H. Saeed, Jinyu Li, Syed Sajid Jan, Vanja Perčulija, Yu Li, Ping Zhu, Songying Ouyang

## Abstract

The current outbreak of Coronavirus Disease 2019 (COVID-19) by a novel betacoronavirus severe acute respiratory syndrome coronavirus 2 (SARS-CoV-2) has aroused great public health concern. Coronavirus has a history of causing epidemics in human and animals. In 2017 an outbreak in piglets by a novel coronavirus was emerged designated as swine acute diarrhea syndrome coronavirus (SADS-CoV) which is originated from the same genus of horseshoe bats (*Rhinolophus*) as Severe Acute Respiratory Syndrome CoV (SARS-CoV) having a broad species tropism. In addition to human cells, it can also infect cell lines from diverse species. Coronavirus host range is determined by its spike glycoprotein (S). Given the importance of S protein in viral entry to cells and host immune responses, here we report the cryo-EM structure of the SADS-CoV S in the prefusion conformation at a resolution of 3.55 Å. Our study reveals that SADS-CoV S structure takes an intra-subunit quaternary packing mode where the NTD and CTD from the same subunit pack together by facing each other. The comparison of NTD and CTD with that of the other four genera suggests the evolutionary process of the SADS-CoV S. Moreover, SADS-CoV S has several characteristic structural features, such as more compact architecture of S trimer, and masking of epitopes by glycan shielding, which may facilitate viral immune evasion. These data provide new insights into the evolutionary relationships of SADS-CoV S and would extend our understanding of structural and functional diversity, which will facilitate to vaccine development.

## Introduction

The risk of major coronavirus (CoV) epidemics in the previous two decades comprising SARS in 2003, Middle East Respiratory Syndrome (MERS) in 2012, SADS in 2017, and the latest 2019 outbreak of human-infecting virulent strain of SARS-CoV-2 have claimed a large number of human lives and livestock. They have shared features of pathogenicity, origin from bats [1–3]. The typical crown-like spike proteins on its surface gives it its name coronavirus. They belong to subfamily *Coronavirinae*, family *Coronaviridae*, order *Nidovirales*, and grouped into four genera including alpha- (α), beta- (β), both CoVs infecting mammals, gamma- (γ) CoVs infecting birds, and delta- (δ) CoVs infecting mammals and birds [4]. The genome of CoV is the second largest of all RNA viruses containing a positive-sense, 27-32 kb in length single-strand RNA (+ssRNA) [5]. CoVs bring about subclinical or respiratory syndrome, central nervous system (CNS) infections, gastrointestinal ailments in humans, and animals [6, 7]. These infections are accountable for 30% of the respiratory illnesses, atypical pneumonia [3, 8], and show a tendency for interspecies transmission, which happened rather often during CoV evolution and shaped the diversity of CoVs [9].

Porcine CoVs are chief health concerns of the pigs that cayses considerable enteric and respiratory infections causing agents of swine. A novel CoV instigating SADS (also called as SeACoV and PEAV) emerged since August 2016. It is associated with HKU2-related *Rhinolophus* bat CoV with death rate up to 90% in piglets of 5 days or younger in Guangdong territory pig breeding farms and accounted for the killing of around 25 000 piglets [10]. SADS-CoV has a position with the class α-CoVs, and it is enveloped +ssRNA virus [2, 6, 11]. The intracellular entry of SADS-CoV relies on a precise interaction amid virion and the host cell. The disease is initiated by the interplay of the viral particle with specific cell surface S trimmer [12].

CoVs from various genera display evident serotypes, for the most part, because of the dissimilarity of their envelope-anchored diversified S proteins. S protein is of around 1300 amino acids, has a place with class I viral combination protein, including SARS-CoVs, human immunodeficiency virus (HIV) envelop glycoprotein 160 (gp160), influenza virus haemagglutinin (HA), paramyxovirus F and Ebola virus glycoprotein [13]. The S protein includes three sections: a large ectodomain, a single-pass transmembrane anchor, and a small intracellular tail. The ectodomain comprises of an N-terminal receptor-binding domain (RBD) viral attachment and entry subunit S1 (approximately 700 amino acids) forming a crown-like structure and a membrane-fusion C-terminal subunit S2 (approximately 600 amino acids) leading to an assembly of architecture, i.e., CTD is sandwiched by its NTD [14]. In addition, CTDs with the assistance of S1 retains a “lying down” inactive state, which transits to “stand up” active state on the S trimer for efficient receptor binding [7, 15–17].

The envelope trimeric S protein is vital for identifying host tropism and transmission limits. It intercedes receptor binding to approach host cell cytosol utilizing acid-dependent proteolytic cleavage of S protein by a cathepsin, TMPRRS2, or additional protease ensued by fusion of the membrane [18, 19]. CoVs utilize an assortment of receptors and triggers to initiate fusion [12]. Receptor binding is fundamental to host-pathogen interaction and pathogenesis. Numerous α-CoVs, for example, human CoV (HCoV-229E), utilize aminopeptidase N (APN) as its receptor, SARS-CoV, HCoV-NL63 and SARS-CoV-2 employ angiotensin-converting enzyme 2 (ACE2), and MERS-CoV ties to dipeptidyl-peptidase 4 (DPP4). However, receptor investigation of SADS-CoV demonstrated that none of the known CoV receptors, i.e., APN, ACE2 and DPP4, is crucial for cell entry that shows the divergence and significance of S protein in SADS-CoV infection [10]. Along these lines, considering the clinical indications of this advancing virus, there is an essential need for primary and practical comprehension of SADS-CoV to expand mechanisms of viral entry and pathogenesis in pigs.

Notwithstanding the significance of the S protein in viral entry and host immunity, high-resolution structural details of this large macromolecular machine have been hard to acquire. In this study, a cryo-EM structure of SADS-CoV S glycoprotein at 3.55 Å resolution is determined. Correspondingly in the present investigation of SADS-CoV, we portrayed the general structure of the S protein and the organization of its structural components. In light of the structures and functions of these essential components, we detail the evolutionary perspective of SADS-CoV S in comparison with S proteins from different CoV genera. In our study, SADS-CoV S forms a “lying down” conformation following the transition from inactive to an active state, and ACE2 is not the receptor of SADS-CoV. The structural alignment suggests that SADS-CoV S is located between α-CoV and β-CoV clade. Hence, our results extend the understanding of critical structural and evolutionary insights into SADS-CoV S, comprising the CTD/RBD essential for receptor recognition and viral entry. This study provides the epidemiological and evolutionary information of this novel CoV in China, which is deteriorating economically important swine industry, and highlights the urgency to develop effective measures to control SADS-CoV.

## Results and discussions

### Overall structure of SADS-CoV spike protein

In order to investigate the role of SADS-CoV S in its invasion, we aimed to solve the high-resolution structure of SADS-CoV S. Ectodomain of SADS-CoV S was expressed, and purified from media of insect cells after 3 days of infection by baculovirus. A fusion peptide of general control protein GCN4 at the C-terminal end of S was used to promote the protein to form a trimer [20] (**Fig. 1A**). The size-exclusion chromatography (SEC) result showed that the protein sample is a trimer in the solution and the purity of more than 95% examined in SDS-PAGE analysis (**Fig. S1A, S1B**). The protein displayed high homogeneity in cryo-EM screening and was then diluted to 0.63 mg ml^−1^ for the data collection. Cryo-EM micrograph movies were collected on a Gatan K2 direct electron detector mounted on an FEI Titan Krios electron microscope (**Fig. S2**). Following reference-free 2D classification of the S protein, we determined a three-dimensional (3D) structure of the SADS-CoV S trimer at 3.55 Å resolution judged by the gold-standard Fourier shell correlation (FSC) criterion of 0.143 (**Fig. 1B, 1C and Table 1**). The resolved atomic structure of prefusion SADS-CoV S ectodomain covers almost all of the key structural elements, as shown in **Fig. 1A** except residues 81-101 of NTD and residues 999-1068 of HR-C (**Fig. 1D**, **Fig. S3**). Forty-five (15 on each subunit) N-linked glycans spread over the surface of the whole S trimer with another 15 predicted but not observed. In addition, the protein trimer is stabilized by 30 pairs (10 on each subunit) of disulfide bonds (**Fig. S4**). The S trimer shows a dome-like shape, with three S1 heads forming a crown-like structure and sitting on top of a trimeric S2 stalk (**Fig. 1D, 1E**, **and Movie S1**). The trimer spike has a length of 143.2 Å from S1 to S2 and a width of 111.3 Å and 61.4 Å at S1 and S2, respectively (**Fig. 1E**). Three S1-CTDs are located at the top center of the S trimer arranged as a small triangle, whereas three S1-NTDs are located on the lower and outer side of S1-CTDs arranged as a big triangle. This architecture, i.e., CTD sandwiched by its own NTD and the adjacent NTD, comes into being one side of the big triangle (**Fig. 1F**). The cleavage site of S1 and S2 subunits locates between residues 544 (Val) and 545 (Arg). The central helicase N (CH-N) and C (CH-C) of S2 from each subunit form a six-helix bundle in the core of S trimer. Heptad repeats N (HR-N) contains four helices connected by three loops locked between the S1 and S2 subunit at the outside of the S2 stalk, while part of the HR-C is missed in the structure (**Fig. 1G**). Each monomeric subunit of S1 contains two major domains, S1-NTD and S1-CTD which are used to bind attachment sialic acid receptors or protein receptors and play a vital role in the CoV entry, and two subdomains, SD1 and SD2 whose function are not very clear (**Fig. 1H-1I**, **Movie S1**).

**Fig. 1.**
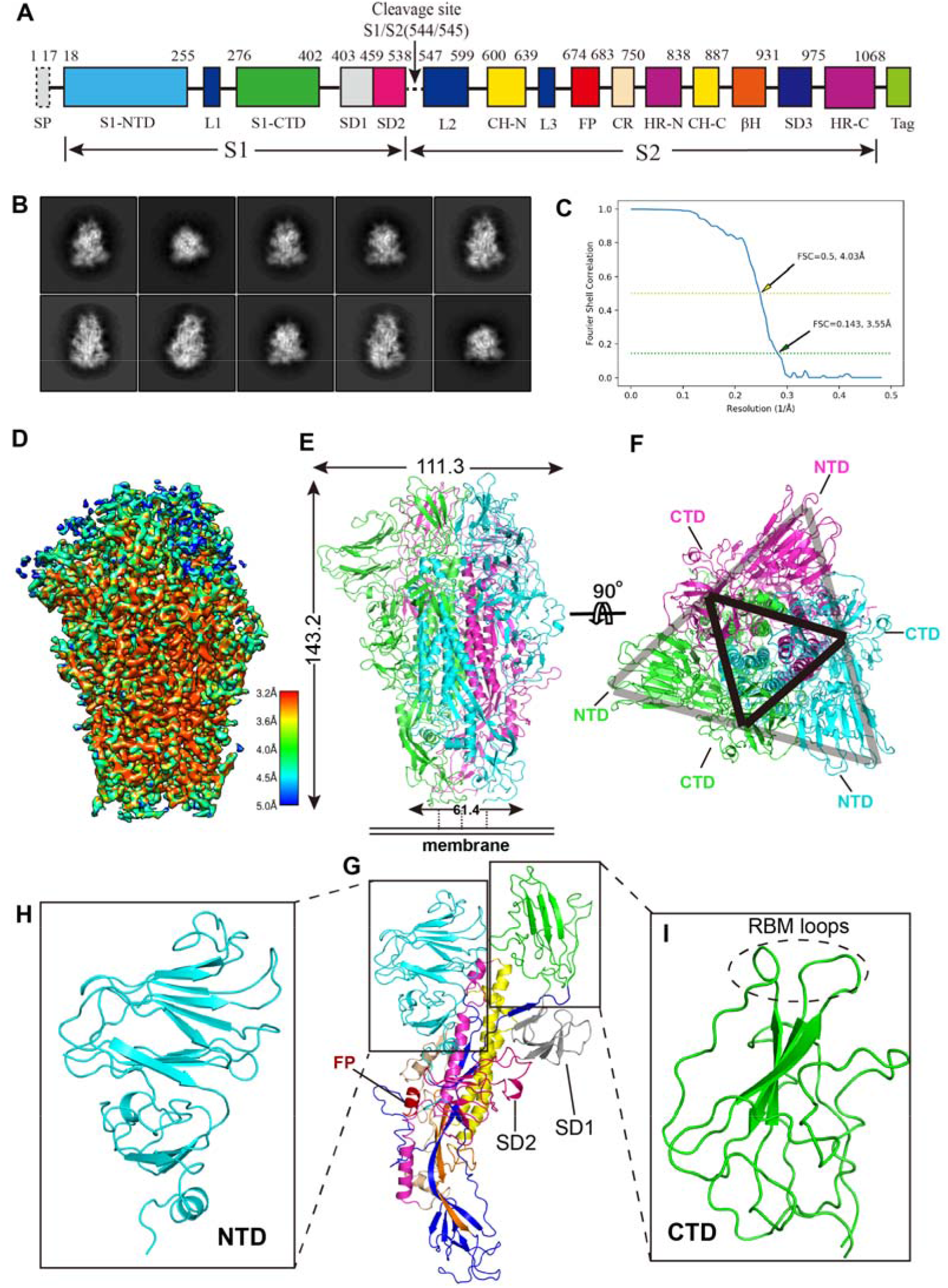
The 3.55 Å cryo-EM structure of SADS-CoV S in the prefusion conformation. (A). Representative 2D class averages in different orientations of SADS-CoV S trimer. (B). Gold-standard FSC curves of SADS-CoV S. The resolution was determined to be 3.55 Å, and a horizontal yellow indicates the 0.5 cut-off value dashed line. (C). Schematic drawing of SADS-CoV S. S1: receptor-binding subunit. S2: membrane-fusion subunit. NTD: N-terminal domain of S1. S1-CTD: C-terminal domain of S1. SD1-3: subdomain 1-3. FP: fusion peptide. CH-N and CH-C: central helices N and C. HR-N and HR-C: heptad repeats N and C. Tag: GCN4 trimerization tag followed by His 6 tag. (D). Final cryo-EM density map of SADS-CoV S colored according to the local resolution. (E). Cryo-EM structure of SADS-CoV S in the prefusion conformation. Each of the monomeric subunits is colored in green, cyan and magenta, respectively. (F). Same as (E), but the dome is rotated 90 degrees to show the top view of the trimer, the CTDs arrange as a small triangle and is indicated by a black triangle. The two adjacent NTDs sandwich one CTD to form one side of a big triangle, which is indicated by a gray triangle. (G). Cryo-EM structure of the SADS-CoV S monomeric subunit. The structural elements are colored in the same way as in panel (A). (H-I). Structures of S1-NTD (cyan) and S1-CTD (green), the putative RBM loops are indicated by a dashed cycle.

**Table 1.** Data collection and refinement statistics.

| Dataset | SADS-CoV S ectodomain |
| --- | --- |
| **Data collection** |  |
| EM equipment | FEI Titan Krios |
| Voltage (kV) | 300 |
| Detector | K2 Summit |
| Pixel size (Å) | 1.04 |
| Electron dose (e<sup>-</sup>/Å<sup>2</sup>) | 60 |
| Defocus range (µm) | -1.8 ~ -2.5 |
| **Reconstruction** |  |
| Software | RELION 3.0 |
| Particle Numbers | 35,000 |
| Symmetry | C3 |
| FSC threshold | 0.143 |
| Final resolution (Å) | 3.55 |
| Map-sharpening *B* factor (Å<sup>2</sup>) | -152 |
| **Model building** |  |
| Software | Coot 0.8.9 |
| **Refinement** |  |
| Software | Phenix-1.18rc1 |
| CC_mask | 0.78 |
| Bond lengths (Å) | 0.007 |
| Bond angles (°) | 1.389 |
| **Validation** |  |
| Clashscore | 5.90 |
| Rotamers outliers (%) | 1.57 |
| **Ramachandran plot** |  |
| Favored (%) | 89.85 |
| Allowed (%) | 8.89 |
| Outliers (%) | 1.26 |

### Structural evolution of coronavirus spike protein

The S protein from all the four different genera of the CoVs packs a crown-like structure by three monomeric subunits, which can be divided into two packing modes: the cross-subunit packing mode and the intra-subunit packing mode [21]. Our SADS-CoV S structure takes an intra-subunit quaternary packing mode where the NTD and CTD from the same subunits pack together by head to head. In SADS-CoV S, the three NTDs from different subunits are located at the vertices of the big triangle formed by the S1 subunits and sandwich the CTDs to the center of the triangle. As a result, the putative receptor-binding moieties located on the top of CTDs are also wrapped at the center of the crown-like trimer (**Fig. 2A**). Accordingly, the geometry of SADS-CoV S resembles the prefusion structures of other α- and δ-genera with the intra-subunit packing mode and is more compact than those of β- and γ-genera that use the cross-subunit packing mode (**Fig. 2A-2E**). Interestingly, although SADS-CoV belongs to the α-genera, its spike protein doesn’t have the domain 0, which is commonly found in other α-genera CoVs, e.g., NL63-CoV and PEDV (**Fig. 2F-G**) [20].

**Fig. 2.**
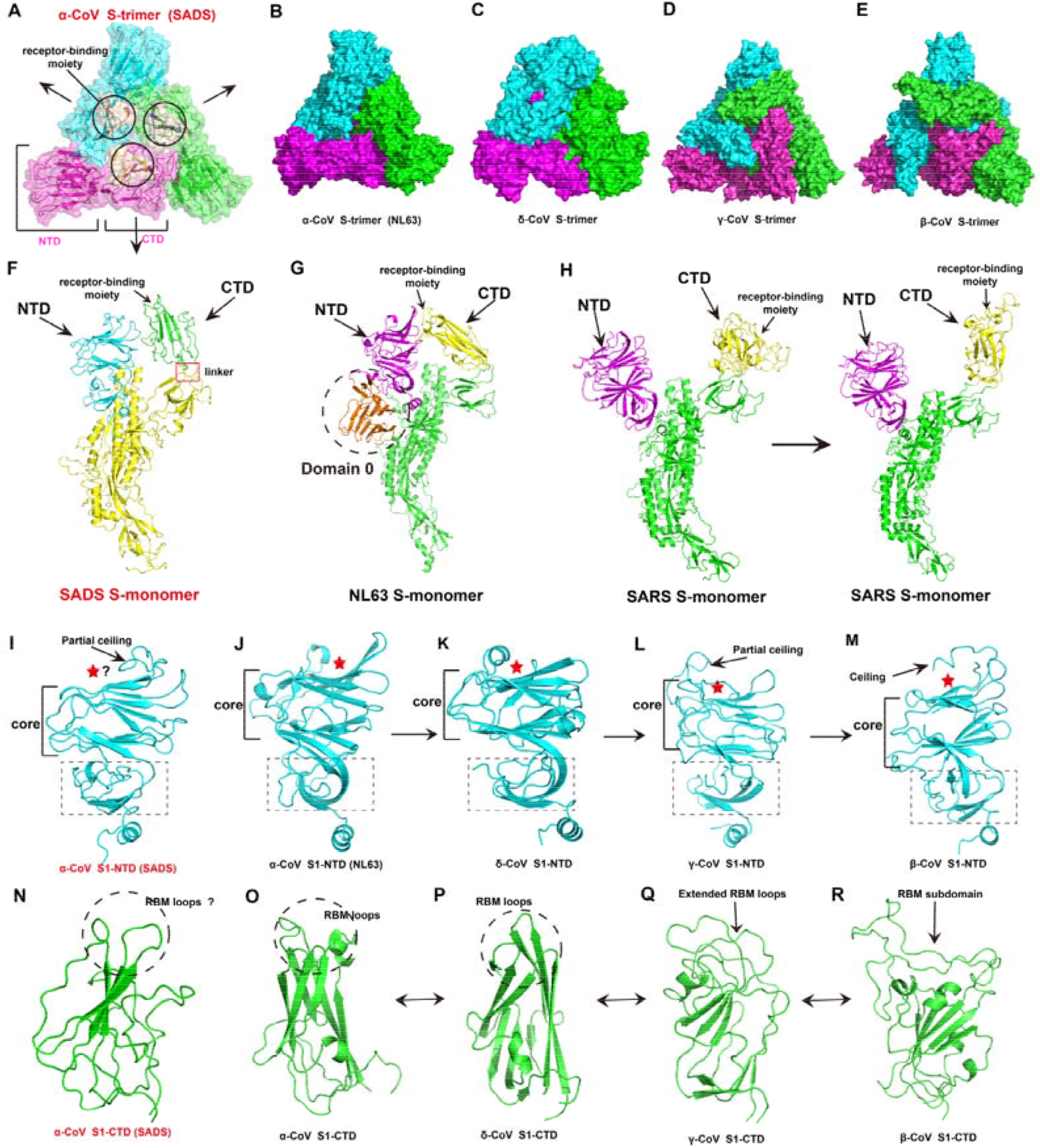
Structural evolution of coronavirus spike protein. (A). The architecture of the SADS-CoV S1 subunit. Three subunits are shown as different colors; one of the NTD and CTD indicated as magenta, the receptor-binding moieties are shown as wheat and indicated by cycles. (B-E). The architecture of the four coronavirus genera S subunit. (B): α-CoVs S-trimer, NL63-CoV, PDB: 5SZS; (C) δ-CoVs S-trimer, PdCoV, PDB: 6BFU; γ-CoVs S-trimer, IBV, PDB: 6CV0; β-CoVs S-trimer, SARS-CoV, PDB: 5X58. (F). Structures of S1 monomer from SADS-CoV. A red frame indicates the linker between the CTD and SD1. The NTD, CTD, and receptor-binding moiety are indicated by a black arrow, respectively. (G). Structures of S1 monomer from NL63-CoV, the additional domain 0 is indicated by a dotted cycle. (H). Structures of S1 monomer from SARS-CoV. The left panel shows the S1 subunit in a “lying down” state, while the right panel shows the S1 subunit in a “stand up” state. The conformational change of the subunits from “lying down” to “stand up” help the trimer to expose receptor-binding moieties and then switch inactive state to active state. (I-M). Structures of NTDs from different genera. The core structures and subdomain (dotted rectangle) of NTDs are labeled, respectively. Partial ceiling and ceiling are indicated by a black arrow. The sugar-binding site or putative sugar-binding site in sugar-binding NTDs are indicated by a red star. (I): SADS-CoV S1-NTD; (J): NL63-CoV S1-NTD (PDB: 5SZS); (K): PdCoV S1-NTD (PDB: 6BFU); (L): IBV S1-NTD (PDB: 6CV0); (M): SARS-CoV S1-NTD (PDB: 5X58). (N-R). Structures of CTDs from different genera. The RBM loops and putative RBM loops are indicated by dotted cycles, while the extended RBM loops and RBM subdomain are indicated by a black arrow. (N): SADS-CoV S1-CTD; (O): NL63-CoV S1-NTD (PDB: 5SZS); (P): PdCoV S1-CTD (PDB: 6BFU); (L): IBV S1-CTD (PDB: 6CV0); (M): SARS-CoV S1-CTD (PDB: 5X58).

Different from the receptor-binding inactive state of these structures whose all three CTDs in “lying down” positions (**Fig. 2A-2E**), the previous study captured several β-CoVs S which contain one or two, even all three CTDs in “stand up” positions [7, 15, 17, 22, 23]. The engagement of the CTDs of SARS S trimer helps it to keep one or more CTDs in the “stand up” position and facilitate the binding of ACE2 or neutralizing antibodies. If all the three CTDs are in the “lying down” position, it is not possible to bind ACE2 due to the partial binding sites are hidden and steric clashes between binding factors. This kind of conformation represents an inactive state (**Fig. 2H**). The CTDs in our structure keep a “lying down” state, and the receptor-binding moieties are partially concealed. It needs to “stand up” on the spike trimer and release the steric clash for efficient receptor binding. The linker between CTDs and S1 subdomains workes as a hinge to facilitate the conformation change of CTDs from “lying down” to “stand up”, furthermore, transit the receptor-binding inactive state to active state (**Fig. 2F**).

It is noteworthy that stronger interactions in the SADS-CoV S trimer may become obstacle conformational change and dissociation of S1-CTD in the prefusion process (**Fig. S5**). Moreover, to probe the intrinsic mobility of the CTD/RBD of S trimer in the apo form (or to see whether the CTD/RBD of S trimer can spontaneously expand to an open state), we performed 2000 ns-long coarse-grained (CG) molecular dynamics (MD) simulations [24]. A Martini CG model was initially generated using the “lying down” state structure (**Fig. S6**). The center-of-mass distances between every two CTDs/RBDs across the simulations (**Fig. S6**) indicated that all the three CTDs/RBDs were stabilized in a “lying down” state similar to the structure observed in this study, i.e. the distance differences calculated from simulations and crystal structure were less than 5 Å. Although we did not find apparent expending of the CTDs/RBDs from a “lying down” state to a “stand up” state during the simulations, it did not rule out the co-existence of the two states in the apo form. Because such a large conformational transition may happen over long time scales (ms to s), far beyond the simulations performed here. Taken together, these simulation results may explain the homogeneity of SADS-CoV S observed in our 3D classification and that the different S1-CTD conformations are invisible in our results (**Fig. 1B**).

For the NTDs, the core structure consists of two six-stranded antiparallel β-sheet layers stacked together, which takes the same galectin fold as human galectins and the NTDs from the other genera. Besides the core structure, NTD of SADS-CoV S1 also has a loop (residues 133-150) formed as a partial ceiling-like structure that resembles the partial ceiling of γ-CoV rather than the other α- and δ-coronavirus which do not have a ceiling-like structure, or the β-CoV which has a reinforced ceiling-like structure (**Fig. 2I-M**). Based on the structural similarity between the NTDs from four different CoVs genera, the sugar-binding site in SADS-CoV S1-NTD might also be located in the pocket formed between the core structure and the partial ceiling (**Fig. 2I**). Compared with other CoVs, the structure of the subdomain under the core domain of SADS-CoV S1-NTD has the same situation as the partial ceiling-like structure (**Fig. 2I-M**). The previous study gave the idea that NTDs from the four genera form an evolutionary spectrum in the order of α-, δ-, γ- and β-genera, with α-CoVs NTD probably being the most ancestral [21]. Our structure has also unexpectedly consistent with this conclusion. Furthermore, we propose that the structural evolution of SADS-CoV S1-NTD is more likely to located between the other α-CoVs, δ-CoVs, and γ-CoVs.

For the CTDs, despite there are dramatic sequential differences between SADS-CoV S1-CTD and other CTDs, all of them share similar structural topology (**Fig. 2N-R**). Unlike the other CTDs of α-CoVs and δ-CoVs which have a more compact core β-sandwich structure containing two β-sheet layers, our CTD resembles the γ- and β-CoVs which just has one layer β-sheet and several α-helix and coil structures with a loose packing (**Fig. 2N-R**). Based on the structural comparison of the CTDs of all the four genera, all the other α-CoVS and δ-CoVs use the loops located on the top of the β-sandwich of CTDs as its receptor binding motif (RBM). As a result, we propose that SADS-CoV also uses these loops on the CTD as its RBM (**Fig. 2N-P**). Moreover, the CTDs of γ-CoVs evolve to get extended loops, and the β-CoVs obtain an insertion domain which contains RBM under the host immune pressure, these receptor-binding moieties also located on the top of core domain (**Fig. 2Q-R**). Previous studies have shown that the core structures are two layers β-sandwiches for α- and δ-CoVs CTDs, weakened β-sandwiches for γ-CoVs CTDs, and single β-sheet layer for β-CoVs CTDs. The RBMs are three short discontinuous loops for α- and δ-CoVs CTDs, two reinforced loops for γ-CoVs CTDs, and a single continuous insertion domain for β-CoVs CTDs. The CTDs from different genera give an evolutionary spectrum, using α- and δ-CoVs CTDs as ancestral, γ-CoVs CTDs as the transitional structure and β-CoVs CTDs the downstream structure, but the evolutionary direction could go either way [21]. Hence, we propose that the SADS-CoV S1-CTD located between the other α-, δ-CoVs, and β-CoVs CTDs on the evolutionary spectrum (**Fig. 2N-R**).

### Structural alignment between the S of SADS-CoV, SARS-CoV and SARS-CoV-2

The phylogenetic analysis based on the whole genome and N gene of eight SADS-CoVs and other CoVs showed that all the sequences of SADS-CoVs clustered with bat coronavirus HKU2 to form a well-defined branch and belong to the α-genera [25]. However, surprisingly/interestingly, the phylogenetic analysis of the S genes showed that the S gene cluster can be divided into two groups: α-CoV-1 and α-CoV-2. All of the SADS-CoVs and bat coronavirus HKU2 belong to α-CoV1 which group with the β-CoVs to form the half evolutionary branch of CoVs, whereas the other α-CoVs cluster to the α-CoV-2 [11, 25]. The structural alignment of the SADS-CoV S trimer by Dali analysis (http://ekhidna2.biocenter.helsinki.fi/dali/) implies that SADS-CoV S has a relative conservation with that of β-CoVs, as reflected by the generally high Z-scores and high RMSD (**Table S1**). In addition, the Dali analysis of CTD showed that SADS-CoV S CTD is relatively poorly conserved with β-CoVs, as reflected by the generally low Z-scores and high RMSD. However, it shares roughly identical core folding with other β-CoVs, i.e., HKU9, SARS-CoV, MERS-CoV, HKU1 and OC43 (**Table S2**). Taken together, these results are consistent with that of the sequence alignment.

Based on the phylogenetic analysis, we compared the CTDs of SADS-CoV, SARS-CoV and SARS-CoV-2 by the sequence and structure. The whole SADS-CoV S ectodomain is only approximately 30% sequence identity when compared with the SARS-CoV and SARS-CoV-2, which obstructs structural alignment between SADS-CoV, SARS-CoV and SARS-CoV-2. Especially there is no significant homologous CoV S CTD found in the PDB when the SADS-CoV S CTD sequence as the inquiry. Interestingly, the SADS-CoV S CTD is smaller (about 130 residues) than other CoV CTD (about 170 residues) (**Fig. 3A**). To further investigate the differences between the S of SADS and other CoVs, and deduce the hypothesis of SADS pathogenic mechanisms, we focused on CTD and performed structural alignments using the complexes of CTD (SARS-CoV-2)-ACE2, CTD (SARS-CoV)-ACE2, CTD (NL63-CoV)-ACE2 (**Fig. 3B-3D**) and the CTDs of SARS-CoV-2, SARS-CoV and SADS-CoV (**Fig. 3E**). Surprisingly, although SARS-CoV-2, SARS-CoV and SADS-CoV CTD do not share significant sequence homology and belong to a different family of CoVs (**Fig. 3A**), they share roughly identical organization and core folding (**Fig. 3E**). Notably, SADS-CoV uses the variant loop region as its RBM, which is consistent with the insertion domain of SARS-CoV and SARS-CoV-2 CTDs that is formed by two β sheets and several loops (**Fig. 3E**). Meanwhile, compared with the β-CoVs SARS-CoV-2 and SARS-CoV, the α-CoV NL63 also use its loops to bind ACE2. Although the loops of SADS-CoV and NL63 CTDs share a structural similarity, SADS-CoV cannot use the ACE2 as its receptor. These features imply that both of the variant sequence and the local structure of these loops are combined to determine the receptor binding specificity.

**Fig. 3.**
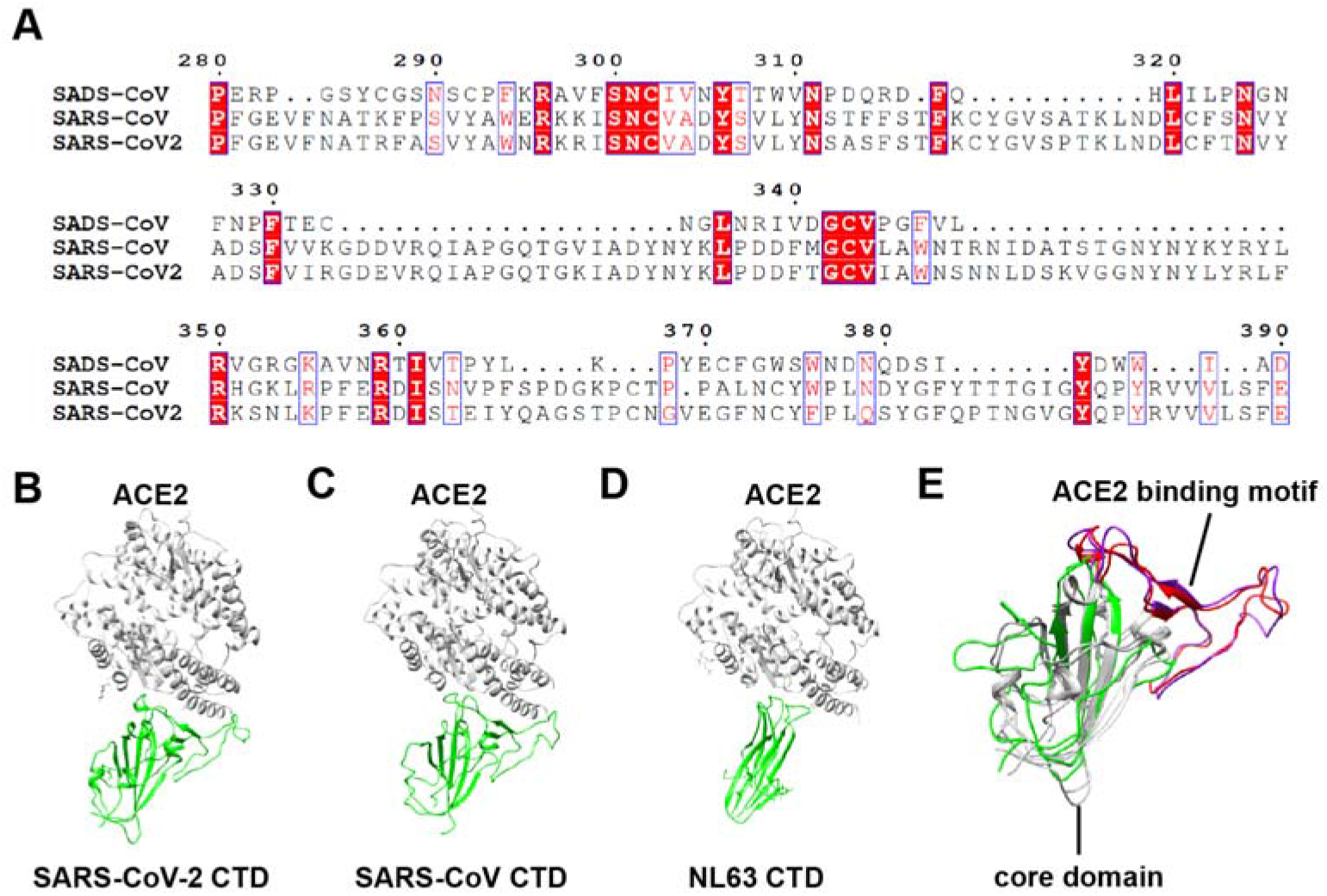
Comparison of the RBD domain of SADS-CoV, SARS-CoV, and SARS-CoV-2. (A). Sequence alignment on CTD of SADS-CoV, SARS-CoV and SARS-CoV-2. (B-D). Structures of SARS-CoV-2 CTD complex with ACE2 (B), SARS-CoV-CTD complex with ACE2 (C), NL63-CoV-CTD complex with ACE2 (D). CTD is colored green and ACE2 is colored gray. (E). Structural alignment of SARS-CoV-2 (colored gray and red), SARS-CoV (colored gray and purple) and SADS-CoV CTD (colored green). Red and purple ribbons indicate the ACE2 binding motif (insertion domain).

### Immune evasion strategies by SADS-CoV S

As a protein located on the surface of the virus, the S proteins mediate viral entry into host cells and also undergo the immune pressure from the immune system of the host at the same time. The structure of SADS-CoV S provides some hints on the immune evasion strategies of SADS-CoV.

On the one hand, the SADS-CoV S has a classical compact structure as other α-CoVs, which uses the intra-subunit packing mode (**Fig. 2A-2B**). As a result, this kind of architecture can maximally reduce the surface area of the S protein to the immune system. Moreover, all of the NTDs and CTDs in a “lay down” station (closed conformation) can further reduce immune pressure (**Fig. 2F**). Nevertheless, the NTDs and CTDs still have the chance to expose for receptor binding. Furthermore, they can also be selected as single- or two-RBD system [4]. Based on the structural alignment of these structures, we speculate that SADS-CoV uses the two-RBD system for which the NTD is used as the attachment receptor-binding domain and the CTD is used as the protein receptor-binding domain. Upon infecting host cells, S1-CTD would need to switch to an open conformation (“stand up”) to render the putative RBM loops accessible to the host receptor. This closed-to-open mechanism can minimize the exposure of the putative RBM loops to the immune system [4].

On the other hand, as one of the immune evasion strategies, masking of epitopes by glycan shielding is usual in CoV evolution [26]. Our map shows that fifty N-linked glycans are spreading over the surface of each S subunit. They are mainly located on the surface of S1 rather than on S2 like HCoV-NL63 (**Fig. 4A**). In contrast to that, HCoV-NL63 S evades host immune surveillance mainly by glycan shielding its S2 epitopes, SADS-CoV spike appears to evade host immune surveillance mainly by glycan shielding its S1 epitopes [26]. In addition, unlike the other α-CoV and δ-CoV whose putative sugar-binding sites are surrounded by glycans, most of the SADS-CoV N-linked glycosylation sites are located on the CTD (**Fig. 4A-4B**). As a result, the putative sugar-binding sites on the NTD are shielded, not by glycans, but by the partial ceiling-like structure on the top of the core structure. As a comparison, this feature is consistent with BCoVS1-NTD [27]. Unlike the HCoV-NL63 receptor-binding residues interacting with domain A belonging to the same protomer, SADS-CoV CTD uses a loop (residues 321-328) to interacting with NTD from the same S1. Both use N-linked glycosylation (residue Asp358) to make them buried and not available to engage the host cell receptor (**Fig. 4C**) [26]. Taken together, SADS-CoV has several unique structural features that may facilitate viral immune evasion.

**Fig. 4.**
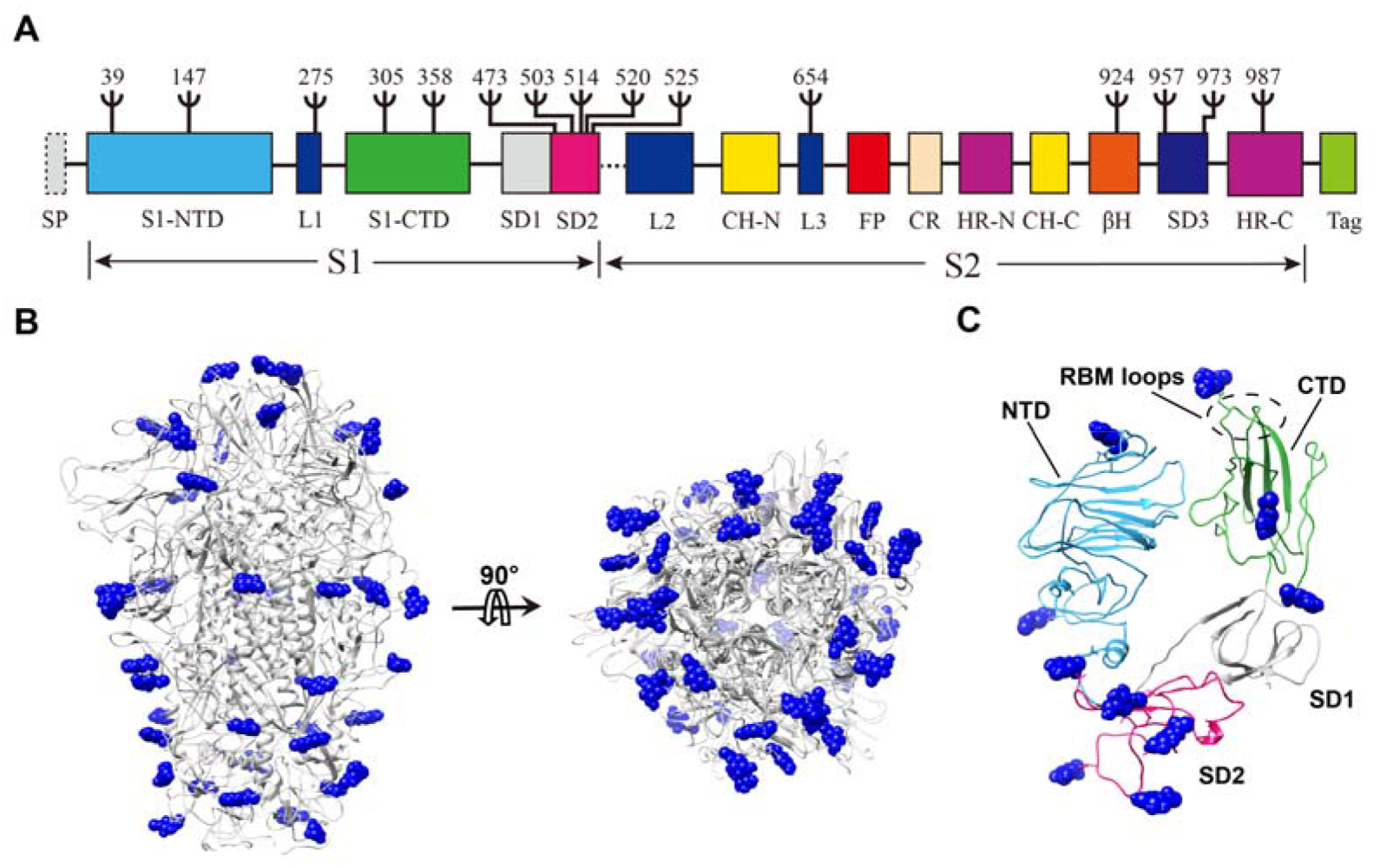
N-linked glycan distribution on the surface of SADS-CoV S. (A). Distribution of observed N-linked glycosylation sites on the one-dimensional structure of SADS-CoV S. Ψ indicates N-linked glycosylate sites. (B). Distribution of observed N-linked glycosylation sites on the three-dimensional structure of the SADS-CoV S. Blue sphere indicates N-linked glycosylate sites. (C). Distribution of observed N-linked glycosylation sites in monomeric S1 (colored as panel A).

## Methods

### Protein expression and purification

SADS-CoV spike glycoprotein (virus strain GDS04; GenBank No.: ASK51717.1) gene was synthesized with codons optimized and inserted into pFastBac vector (Life Technologies Inc.). The ectodomain of SADS-CoV spike protein without the transmembrane anchor and intracellular tail (residues 18-1068) was expressed by the Bac-to-Bac insect cell system (Invitrogen). To get the trimer ectodomain protein, we added a GCN4 trimerization tag followed by a TEV cleavage site and an 8×His-tag at the C terminal of the S protein. The cells were harvested by centrifugation at 4,000×g and remove the cells. Then the supernatant was loaded to Ni-NTA (Nitrilotriacetic acid) resin (Invitrogen) affinity-purified by the C-terminal 8×His-tag. Spike protein was finally purified using Superose 6 HR10/300 column (GE Healthcare) pre-equilibrated with buffer containing 20 mM HEPES (pH 7.5) and 150 mM NaCl, 1 mM DTT and concentrated with a centrifugal filter (Amicon Ultra) to approximately 1 mg/ml and divided into aliquots, flash-frozen in liquid nitrogen.

### Cryo-EM sample preparation and data acquisition

Purified S protein was diluted to 0.63 mg ml-1 with buffer containing 20 mM HEPES pH 7.5, 100 mM NaCl and 2 mM DTT. A 4 μl volume of sample was applied to a glow-discharged Quantifoil copper grid and vitrified by plunge freezing in liquid ethane using a Vitrobot Mark with a blotting time of 3 s. Data collection was performed on a Titan Krios microscope operated at 300 kV and equipped with a field emission gun, a Gatan GIF Quantum energy filter and a Gatan K2 Summit direct electron camera in super-resolution mode, at CBI-IBP. The calibrated magnification was 130,000× in EF TEM mode, corresponding to a pixel size of 1.04 Å. The automated software SerialEM was used to collect 1,000 movies at a defocus range of between 1.8 and 2.3 μm. Each exposure (10 s exposure time) comprised 32 sub-frames amounting to a total dose of 60 electrons Å-2 s^−1^.

### Image processing

Micrograph movie stacks were corrected for beam-induced motion using MotionCor2 [28]. The contrast transfer function parameters for each dose weighting image were determined with Gctf [29]. Particles were initially auto picked with Gautomatch without template and extracted with a 256-pixel by 256-pixel box. Reference-free 2D-class average was performed using RELION [30], and the well-resolved 2D averages were subjected to another iteration particle auto picking as a template with Gautomatch. After iterative 2D-class average in RELION, only particles with best-resolved 2D averages were selected for initial model generation and 3D classification using RELION. The classes with identical detailed features were merged for further auto-refinement with a sphere mask, and post-processed with 3-pixel extension and 3-pixel fall-off around the entire molecule, to produce the final density map with an overall resolution of 3.55 Å. Chimera and PyMOL (https://pymol.org/) were used for graphical visualization [31].

### Model building

*Ab initio* modeling of the spike protein was performed in Coot [32], using structure predictions calculated by Phyre2 [33], the partial structure modeled by EMBuilder [34], and the reference model (PDB: 5X58). Map refinement was carried out using Phenix.real_space_refine [35], with secondary structure and Ramachandran restraints. Cryo-EM data collection, refinement and validation statistics were listed in Table 1.

## Supporting information

Movie for Main text

## ACCESSION NUMBER

The Cryo-EM structure of SADS-CoV S has been deposited into EMDB with accession numbers of EMD-30071. Coordinates and structure factors have been deposited in the Protein Data Bank (PDB) under accession number 6M39.

## Acknowledgements

S.O was funded by National Natural Science Foundation of China grants (31770948, and 31570875) (S.O), The high-level personnel introduction grant of Fujian Normal University (Z0210509), Special Funds of the Central Government Guiding Local Science and Technology Development (2017L3009); P.Z was funded by National Natural Science Foundation of China grants (31425007), grants from the Chinese Ministry of Science and Technology (2017YFA0504700), Strategic Priority Research Program from Chinese Academy of Sciences (XDB08010100); H.G was funded by National Natural Science Foundation of China grants (31900879), Natural Science Foundation of Fujian province grants (2019J05064).

## AUTHOR CONTRIBUTIONS

**Fig. S1.**
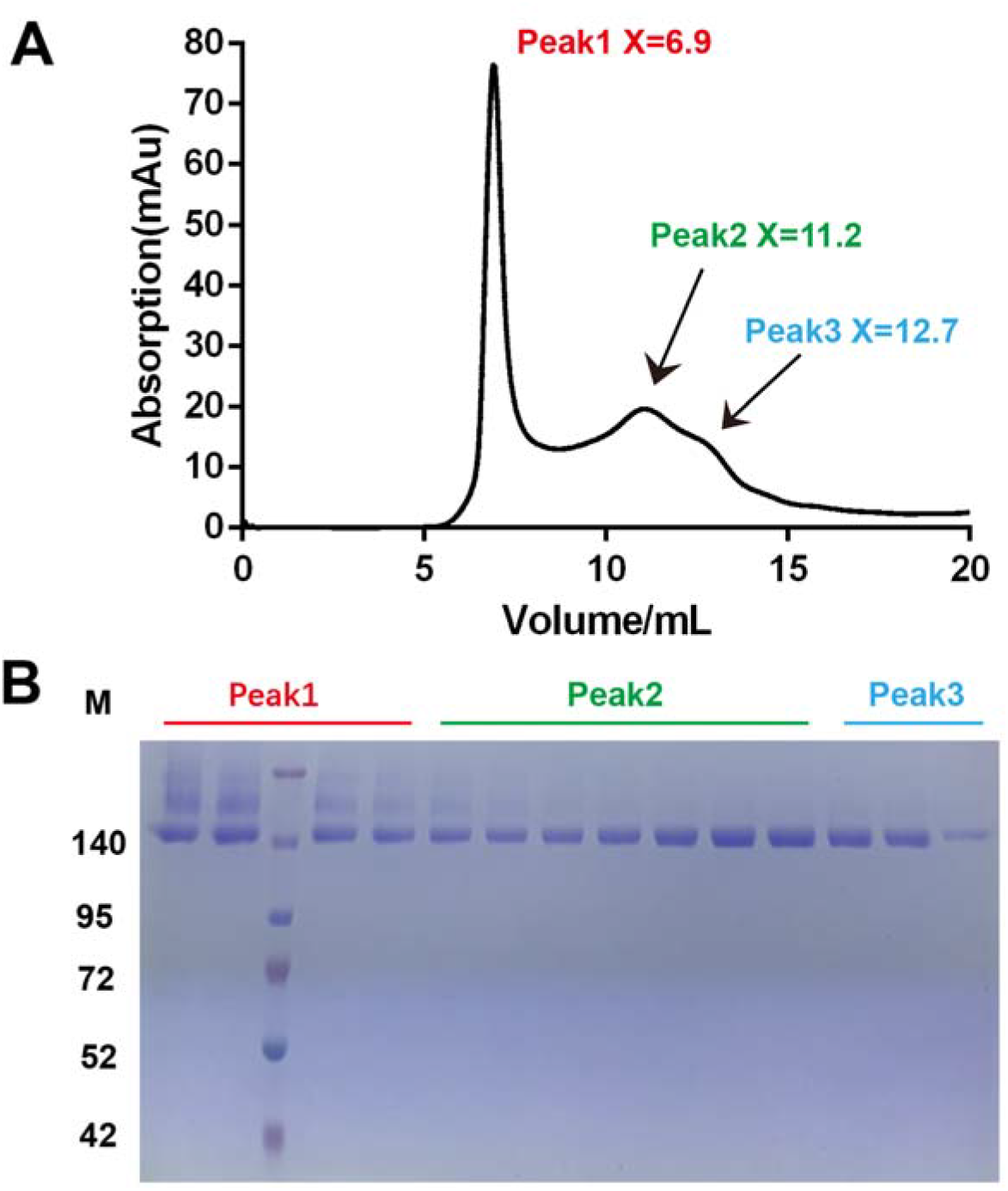
Expression and purification of SADS-CoV S. A. The size-exclusion chromatogram (SEC) of SADS-CoV S gives three peaks and the elution volumes of which are indicated by red, green and cyan, respectively. Data from a Superose 6 10/300 column are shown in the black line. B. The samples from different peaks were checked by SDS-PAGE analysis and the sample from peak 2 (green) is used to analyze further.

**Fig. S2.**
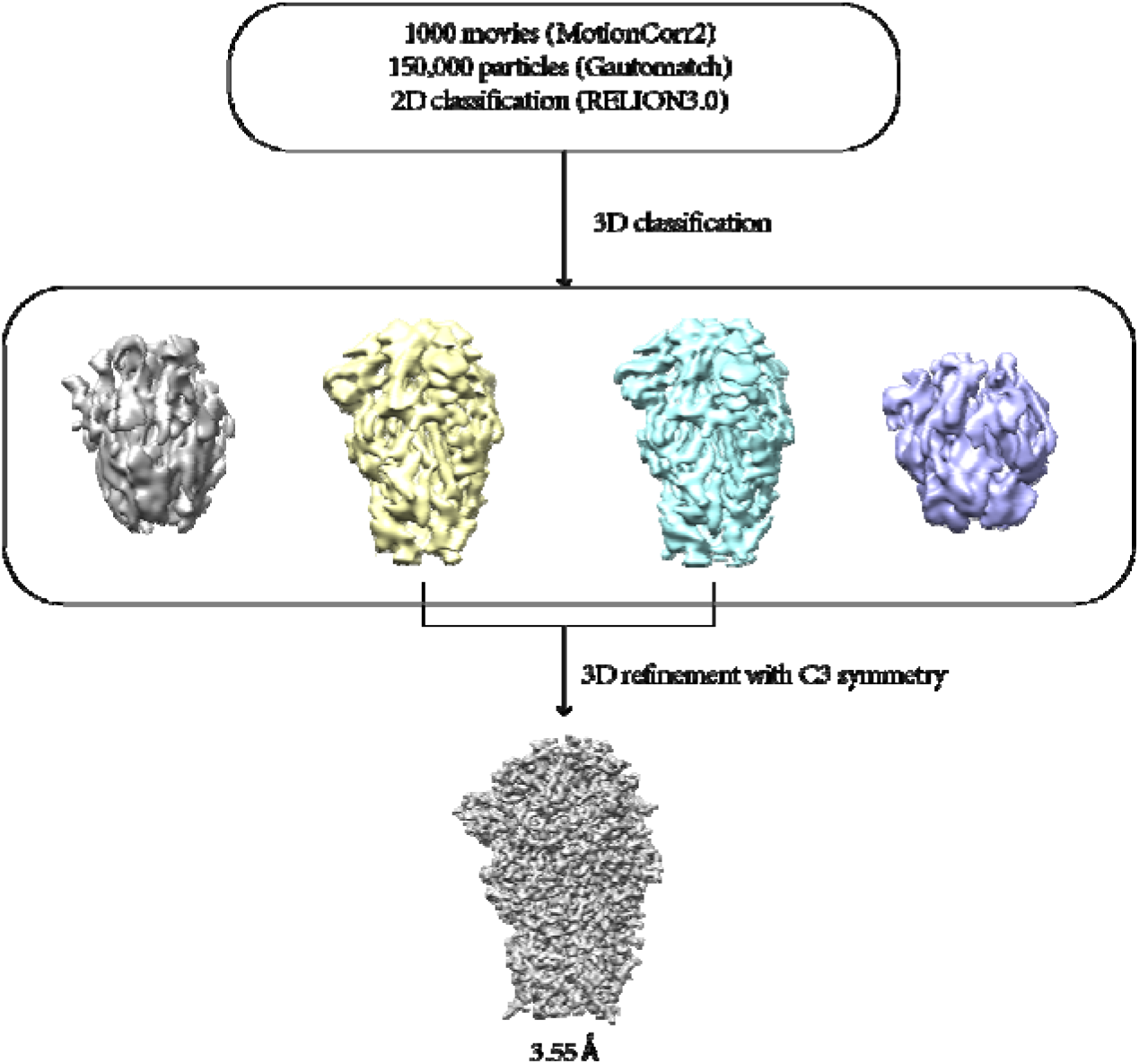
Cryo-EM data processing flow chart of SADS-CoV S.

**Fig. S3.**
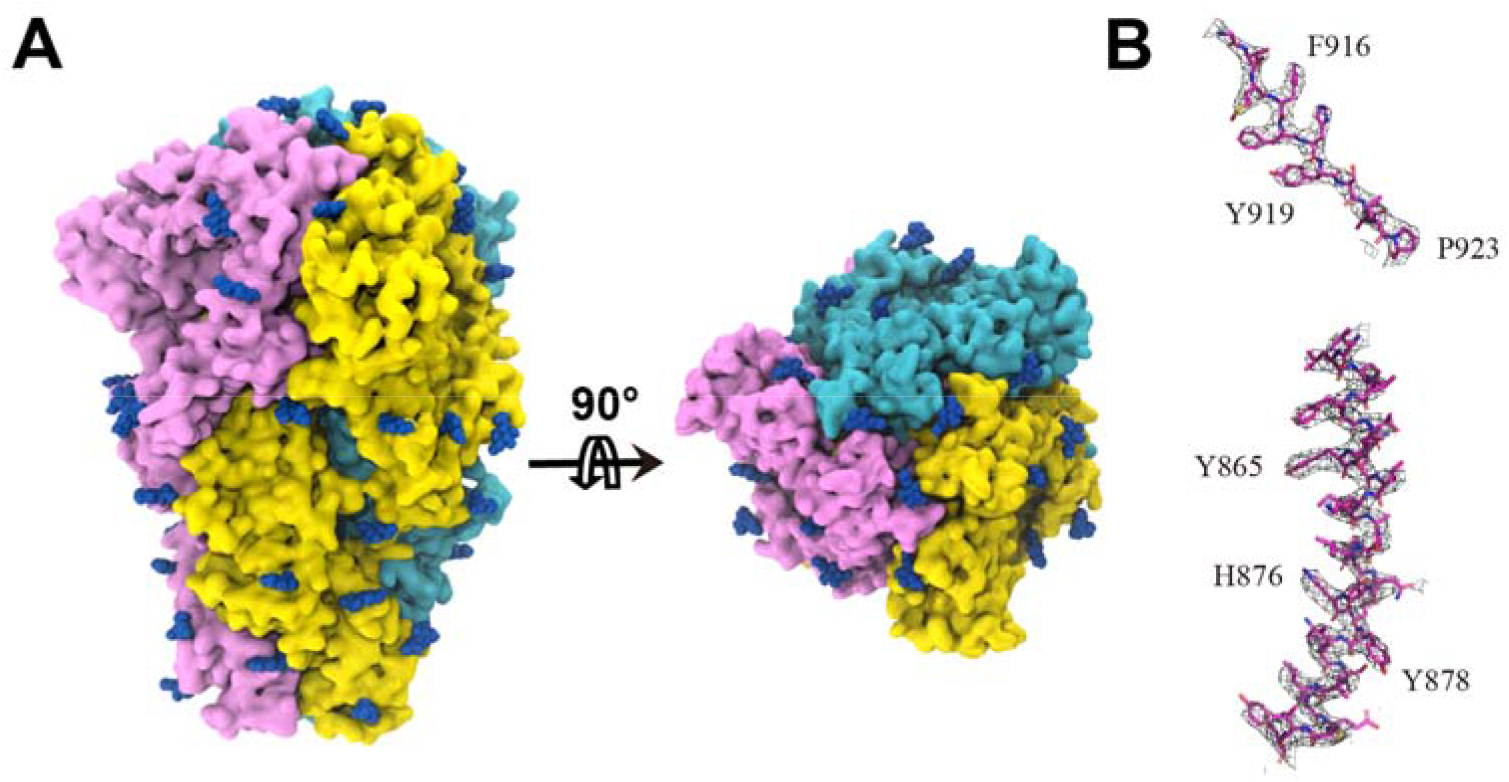
SADS-CoV S structure cryo-EM map (A) and representative density (B). The map is colored by protomer and blue density indicates observed glycosylation sites.

**Fig. S4.**
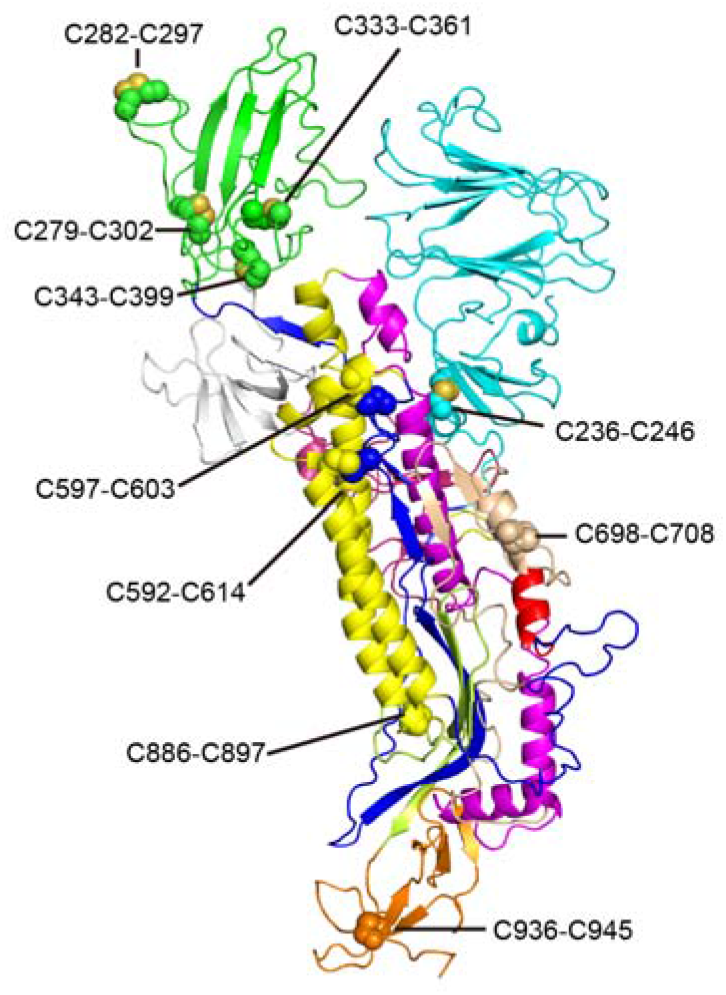
The distribution of 10 pairs disulfide bond on each subunit. (236-246, 287-292, 279-302, 343-399, 333-361, 592-614, 597-603, 698-708, 886-897, 936-945).

**Fig. S5.**
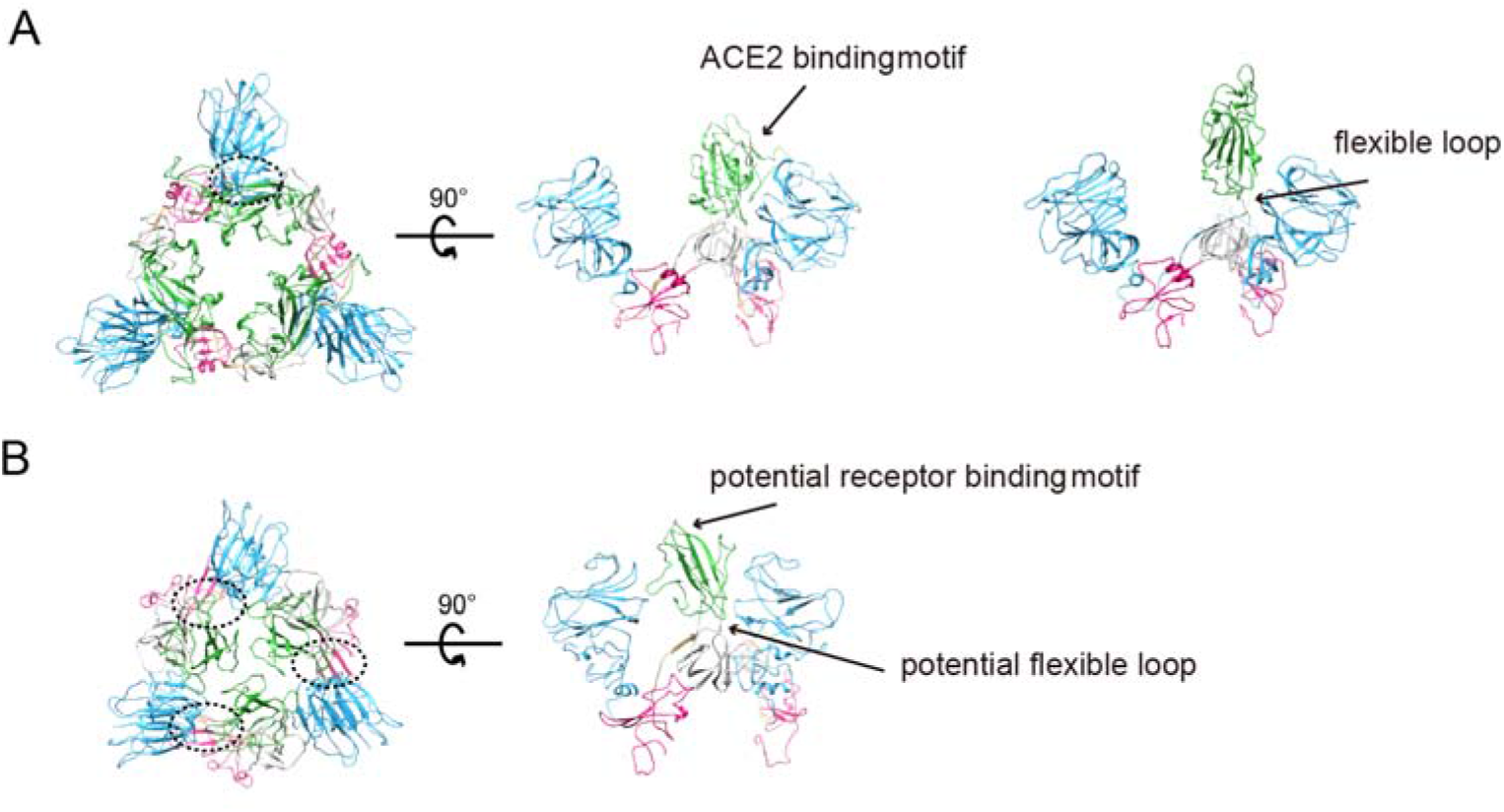
Structural comparison between SARS-CoV spike S1 domain (A) and SADS-CoV spike S1 domain (B). S1-NTD, S1-CTD, SD1 and SD2 are colored by cyan, green, gray and red, respectively. In the native state, only interactions between S1-CTD and S1-NTD of neighbor monomer are visible in SARS-CoV S (dotted circle in panel A). S1-CTD is dissociated from S1-trimer when binding ACE2 in the prefusion state. While in SADS-CoV spike, three interaction regions are visible in the native state (dotted circles in panel B), including S1-CTD and S1-NTD of neighbor monomer, internal S1-CTD and S1-NTD, S1-CTD trimer.

**Fig. S6.**
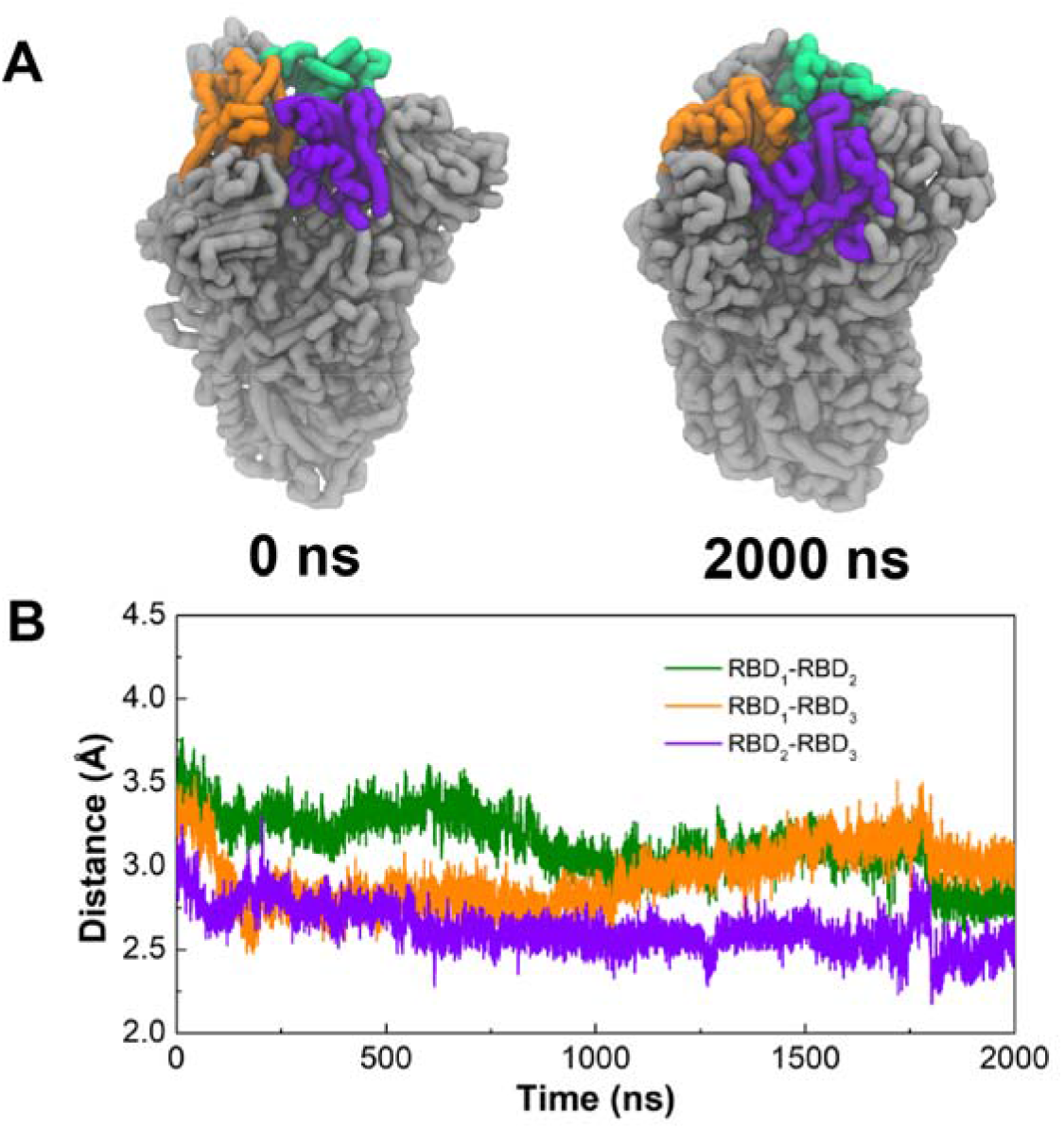
Molecular dynamics of simulations on SADS-CoV S trimer. (A) The initial and final coarse-grained models of SADS-CoV S trimer from the simulations. The three RBDs were highlighted in green, orange and purple, respectively. (B) The center-of-mass distances between every two RBDs in the simulations.

**Table S1.**
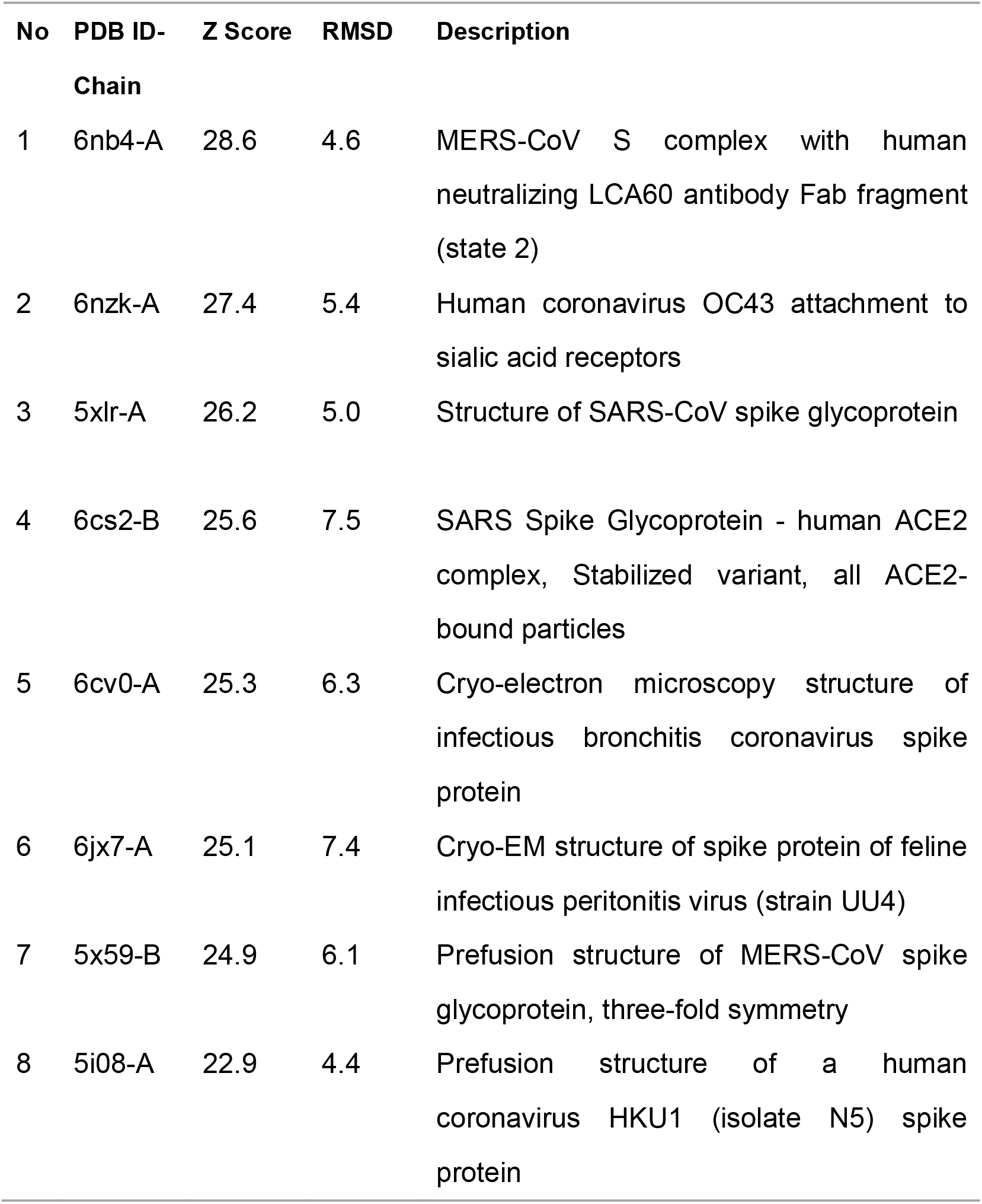
The results of the SADS-CoV S structure comparison by Dali service.

| No | PDB ID-Chain | Z Score | RMSD | Description |
| --- | --- | --- | --- | --- |
| 1 | 6nb4-A | 28.6 | 4.6 | MERS-CoV S complex with human neutralizing LCA60 antibody Fab fragment (state 2) |
| 2 | 6nzk-A | 27.4 | 5.4 | Human coronavirus OC43 attachment to sialic acid receptors |
| 3 | 5xlr-A | 26.2 | 5.0 | Structure of SARS-CoV spike glycoprotein |
| 4 | 6cs2-B | 25.6 | 7.5 | SARS Spike Glycoprotein - human ACE2 complex, Stabilized variant, all ACE2-bound particles |
| 5 | 6cv0-A | 25.3 | 6.3 | Cryo-electron microscopy structure of infectious bronchitis coronavirus spike protein |
| 6 | 6jx7-A | 25.1 | 7.4 | Cryo-EM structure of spike protein of feline infectious peritonitis virus (strain UU4) |
| 7 | 5x59-B | 24.9 | 6.1 | Prefusion structure of MERS-CoV spike glycoprotein, three-fold symmetry |
| 8 | 5i08-A | 22.9 | 4.4 | Prefusion structure of a human coronavirus HKU1 (isolate N5) spike protein |

**Table S2.**
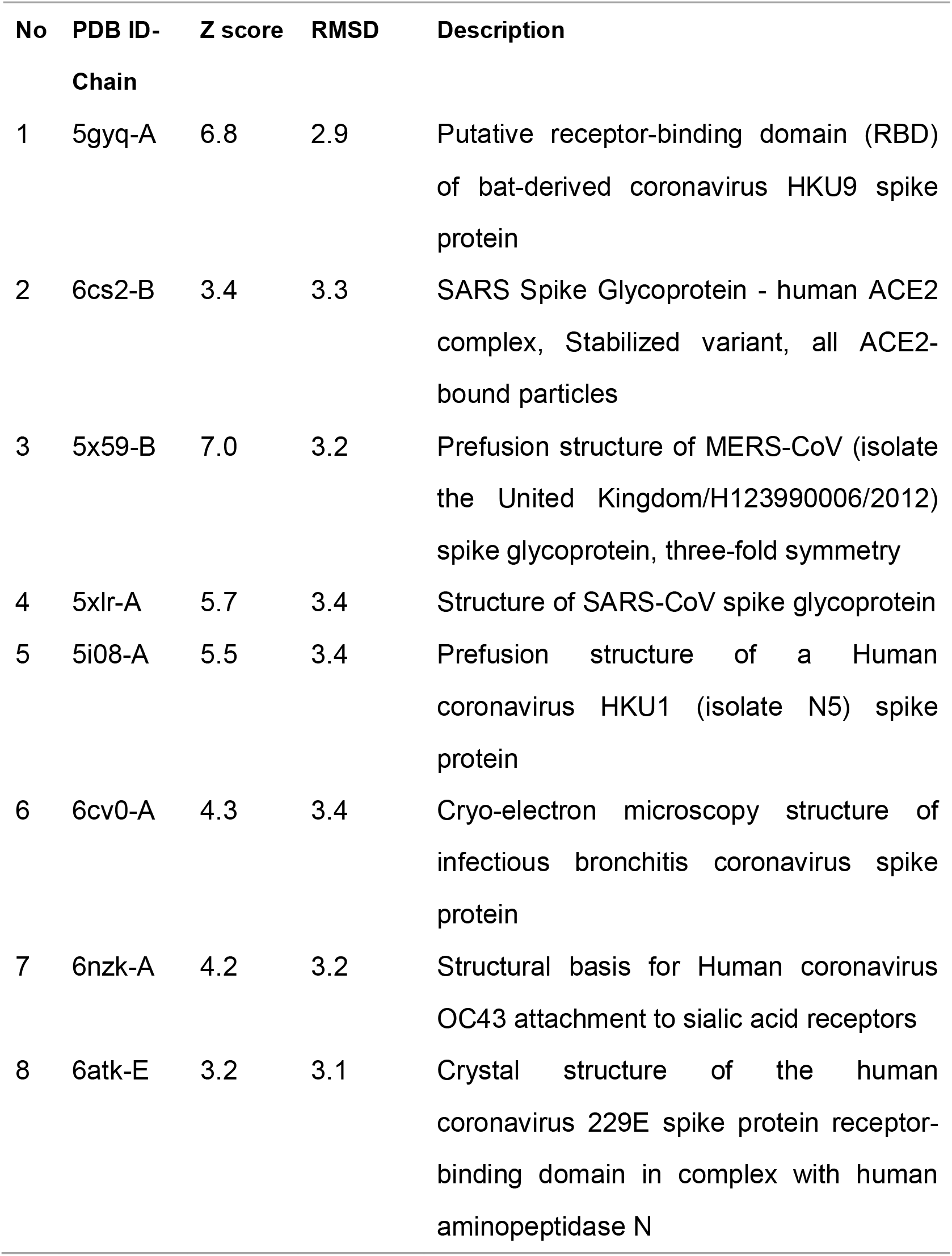
The results of the SADS-CoV S RBD structure comparison by Dali service.

| No | PDB ID-Chain | Z score | RMSD | Description |
| --- | --- | --- | --- | --- |
| 1 | 5gyq-A | 6.8 | 2.9 | Putative receptor-binding domain (RBD) of bat-derived coronavirus HKU9 spike protein |
| 2 | 6cs2-B | 3.4 | 3.3 | SARS Spike Glycoprotein - human ACE2 complex, Stabilized variant, all ACE2-bound particles |
| 3 | 5x59-B | 7.0 | 3.2 | Prefusion structure of MERS-CoV (isolate the United Kingdom/H123990006/2012) spike glycoprotein, three-fold symmetry |
| 4 | 5xlr-A | 5.7 | 3.4 | Structure of SARS-CoV spike glycoprotein |
| 5 | 5i08-A | 5.5 | 3.4 | Prefusion structure of a Human coronavirus HKU1 (isolate N5) spike protein |
| 6 | 6cv0-A | 4.3 | 3.4 | Cryo-electron microscopy structure of infectious bronchitis coronavirus spike protein |
| 7 | 6nzk-A | 4.2 | 3.2 | Structural basis for Human coronavirus OC43 attachment to sialic acid receptors |
| 8 | 6atk-E | 3.2 | 3.1 | Crystal structure of the human coronavirus 229E spike protein receptor-binding domain in complex with human aminopeptidase N |

